# Paraspeckle Protein NONO Regulates Active Chromatin by Allosterically Stimulating NSD1

**DOI:** 10.1101/2025.01.21.634125

**Authors:** Chen-I Hsu, Shenglin Mei, Justin Demmerle, Sarah Ruttenberg, Mahnue Sahn, Jia-Ray Yu

## Abstract

NSD1 is a key lysine methyltransferase for di-methylation of lysine 36 of histone H3 (H3K36me2) essential for the establishment of active chromatin domains. While the loss of NSD1 catalytic activity halts embryonic development and a gain of that drives oncogenesis in leukemia and glioma, the regulatory mechanisms that control NSD1 activity in these processes remain poorly understood. Here, we uncover that NSD1 requires allosteric activation through the aromatic pocket of its PWWP2 domain. Surprisingly, we identify that NSD1-PWWP2 binds to a non-canonical target, nuclear paraspeckle protein NONO, and this protein-protein interaction allosterically stimulates the catalytic activity of NSD1. Mouse embryonic stem cells (mESC) engineered with mutations in the aromatic pocket of NSD1-PWWP2 cannot differentiate into neural progenitor cells (NPC), and genetic depletion of NONO partially phenocopied this defect at cellular and transcriptional levels, potentially explaining the neurodevelopmental disorder phenotypes in NSD1- or NONO-deficient patients. Our work revealed a novel mechanism driving active chromatin domain formation and has critical implications in the interplay between nuclear paraspeckles and active chromatin, and a vulnerability of NSD1 for therapeutic interventions.

**GRAPHIC ABSTRACT:** 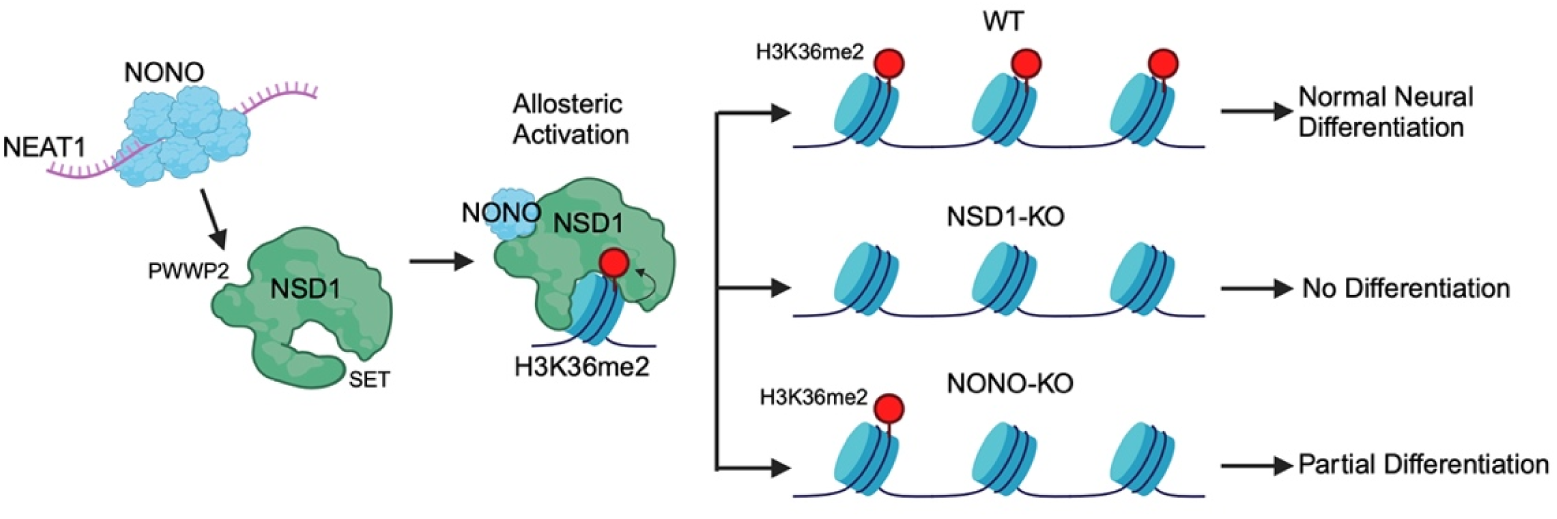

**HIGHLIGHTS:**

1. The catalytic activity of NSD1 requires the aromatic pocket of its PWWP2 domain
2. The aromatic pocket of NSD1-PWWP2 binds to a non-canonical target, paraspeckle protein NONO, which allosterically stimulates NSD1
3. Disruption of the assemblies of paraspeckles dampens global H3K36me2 levels
4. NONO deficiency partially phenocopies the defects of neural differentiation in NSD1-knockout embryonic stem cells

## INTRODUCTION

Mammalian chromatin is a highly compartmentalized structure and a balance between active and repressive chromatin domains is essential for maintaining proper gene expression^1^. Post-translational modifications (PTM) of histones play key roles in this process to ensure normal cell differentiation and animal development^1^. Many mutations in histone-modifying enzymes have been demonstrated to drive human developmental disorders as well as oncogenesis in numerous types of cancer^1,2^. Therefore, how these enzymes are normally regulated in development and dysregulated in pathogenic conditions poses key fundamental questions in chromatin biology. Prior efforts have gained significant insights into understanding how the catalytic activities of these enzymes are orchestrated to establish repressive chromatin domains. For example, it was shown that Polycomb Repressive Complex 2 (PRC2) is allosterically activated through binding to its own catalytic product, tri-methylation of lysine 27 of histone H3 (H3K27me3)^3,4^. This “write-and-read” mechanism constitutes the molecular basis of how repressive chromatin is established and inherited through this self-sustaining, feedback mechanism^3,4^.

While advances have been made to understand the mechanisms of how repressive chromatin is established, little is known about those of active chromatin. Previous work including ours reported that while H3K27me3 and repressive chromatin are compromised, di-methylation at Lysine 36 of histone H3 (H3K36me2) appears to be the most elevated, active histone PTM in diverse types of mammalian cells^5–7^. H3K36me2 structurally hinders the catalysis of H3K27me3^8,9^. In addition to antagonizing H3K27me3 and repressive chromatin, recent work including ours have reported that H3K36me2 plays at least two other key roles in stabilizing active chromatin by 1) recruiting the downstream “reader” proteins LEDGF and HDGF2 that exhibit chaperone-like activities to promote nucleosome disassembly and reassembly to facilitate RNA-Pol II-dependent transcriptional elongation, and 2) recruiting DNMT3A and DNMT3B to shape intergenic and genic DNA methylation patterns, respectively, to prevent undesired transcriptional initiation within active chromatin^10,11^.

In mammals, there are five H3K36 methyltransferases, Nuclear Receptor Binding Set Domain Proteins 1/2/3 (NSD1/2/3), Ash1-like Histone Lysine Methyltransferase (ASH1L), and SET Domain Containing 2 (SETD2)^12^. SETD2 mostly catalyzes H3K36me3, largely independent of the catalysis of H3K36me2^7,13^. Notably, the deposition of H3K36me3 is restricted to gene bodies and has been implicated in regulations of transcription fidelity, RNA splicing, and DNA repair^14^. In contrast, H3K36me2 broadly decorates both genic and large intergenic regions to set the global boundaries of active chromatin^11,13^. Among H3K36me2 methyltransferases, ASH1L appears to be restricted to specific loci, and NSD3 is a weak methyltransferase of the NSD family members, leaving NSD1 and NSD2 as the major enzymatic “writers” for global H3K36me2 in most cell types^15^. In mouse and human genetics, either NSD1 or NSD2 genetic knockout is lethal in mice, and heterozygous mutations in NSD1 or NSD2 drive developmental disorders in human patients^16–19^. In particular, NSD1 is the dominant paralogue in embryonic stem cells (ESC)^20^. NSD1 is dispensable for ESC self-renewal in defined culture conditions but required for ESC differentiation into the three germ layers^20,21^. In human cancer, NSD1 is a cancer dependency gene in Diffuse Midline Glioma (DMG) and has also been reported as an oncogene with its N-terminus truncated to form a gene fusion with nuclear pore protein NUP98^7,22^. However, no clinical grade orthosteric inhibitors are available. Previous biochemical and structural analyses have mainly focused on the catalytic domain of NSD1, but how NSD1 is allosterically or temporo-spatially regulated to license the catalytic process remains largely unknown. A key hurdle is that NSD1 is a very large protein (>300kD) with nearly 50% unstructured/disordered regions, making it difficult to manipulate experimentally. These issues prompted us to establish a novel platform for biochemical reconstitution of NSD1 and investigate the regulatory mechanisms underlying the establishment of H3K36me2-decorated active chromatin by NSD1.

## RESULTS

### Identification of the PWWP2 domain as being essential for NSD1 catalytic activity but independent of its chromatin binding

To investigate how NSD1 is regulated, we screened possible functional domains independently of its catalytic module. We reasoned that since NSD1 is associated with active chromatin, it may bind to nascent RNA transcripts which may in turn regulate its activity. To test this hypothesis, we examined NSD1-RNA interactions using the Proteomic Identification of RNA-Binding Regions (RBR-ID) database and targeted five main RNA-binding domains (RBDs) of NSD1, including the RBD1, 2/3, 4, and PWWP1 (**Figure S1A**)^23^. Besides PWWP1, other RBDs reside within the large unstructured/disordered region of NSD1 (**Figure 1A**). Although the PWWP2 domain does not bind to RNA, the necessity of PWWP2 alone has not been tested before and therefore we included PWWP2 in our domain screening (**Figure S1A**). To test how these domains regulate NSD1 catalytic activity, we generated a NSD1/2 double knockout (dKO) cell line of HEK293T to deplete global H3K36me2 levels and then rescued NSD1/2-dKO cells with either wildtype (WT) or mutant NSD1 using a PiggyBac transposon-based system (**Figure S1B**). We found that single deletion of RBDs or PWWP1 alone does not impact NSD1 activity to catalyze H3K36me2 but surprisingly, the deletion of PWWP2 (NSD1^ΔPWWP2^) completely abrogated NSD1 catalytic activity (**Figure 1B and S1C**). As other mammalian PWWP domains have been shown to regulate protein-chromatin interactions, we asked if the loss of PWWP2 diminished either the intrinsic catalytic activity of NSD1 or the binding of NSD1 to chromatin for access to nucleosomal substrates^10,11,24^. To investigate these possibilities, we performed Chromatin Immunoprecipitation followed by deep sequencing (ChIP-seq) comparing WT and NSD1^ΔPWWP2^ rescued HEK293T cells. ChIP-seq signals were normalized by spike-in Drosophila chromatin. Intriguingly, while NSD1^ΔPWWP2^ had little catalytic activity for H3K36me2, it exhibited much stronger binding to chromatin than NSD1-WT (**Figure 1C and S1D**). This observation suggested that NSD1^ΔPWWP2^ is an inactive trapping mutant bound to its nucleosomal substrates without completing the catalysis of H3K36me2 and releasing from chromatin.

**Figure 1.**
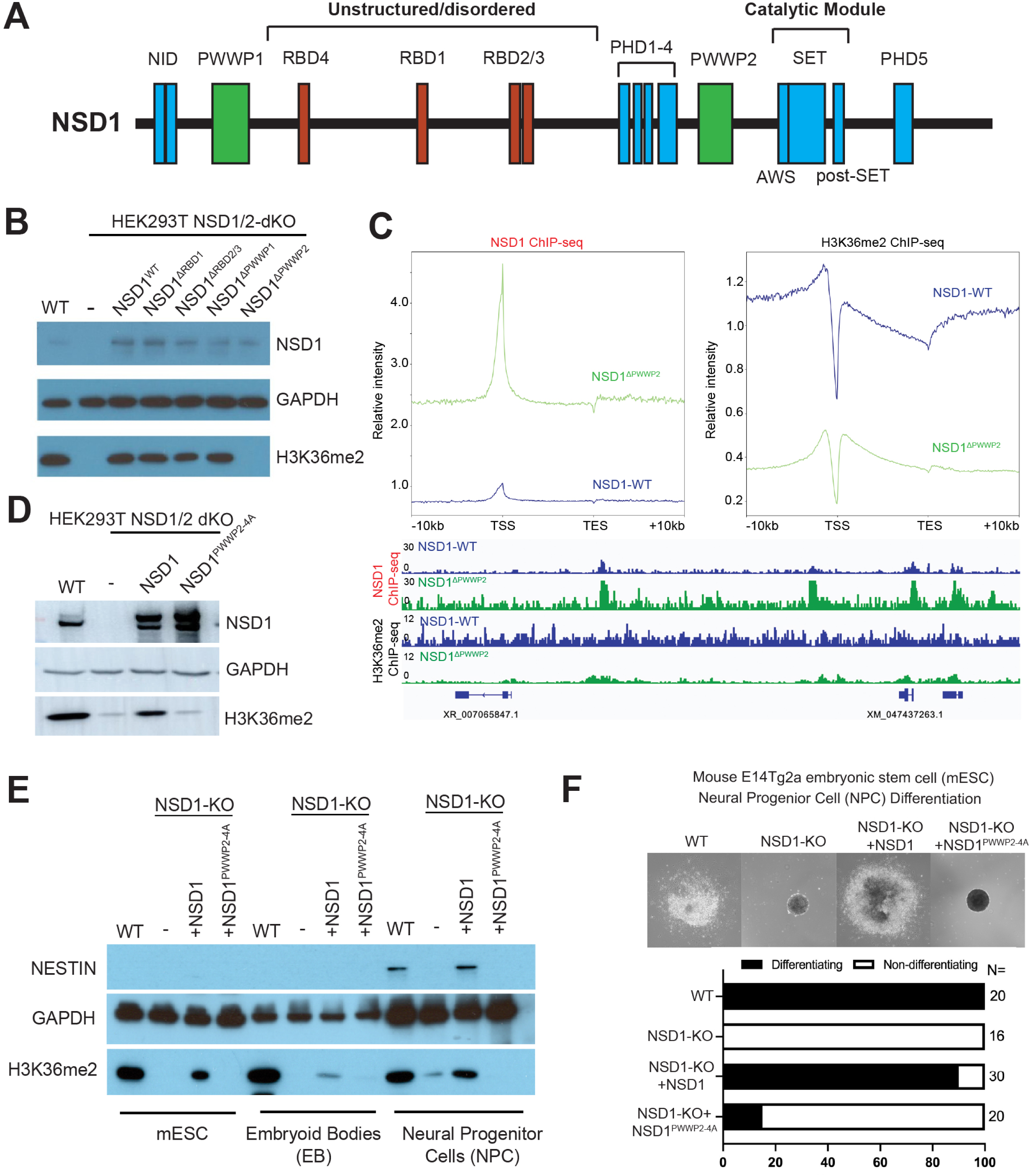
The aromatic pocket of NSD-PWWP2 is essential for NSD1’s catalytic activity and early neural differentiation but dispensable for NSD1’s recruitment to chromatin. (A) Schematic illustration of annotated and putative functional domains of NSD1. NID, nuclear receptor interaction domain; PWWP, Pro-Trp-Trp-Pro domain; RBD, RNA-binding domain; PHD, plant homeodomain; AWS, associated with SET domain (also refereed as pre-SET); SET, su(var), enhancer of zeste, trithorax domain. (B) Western blot of NSD1, GAPDH, and H3K36me2 for WT or NSD1/2-dKO HEK293T cells stably rescued with WT or NSD1 mutants. (C) Top, meta-analysis profiling of genome-wide NSD1 and H3K36me2 ChIP-seq signals within −10kb of TSS to +10kb of TES. Bottom, representative track images of NSD1-WT or NSD1^ΔPWWP2^ in HEK293T NSD1/2-dKO rescued background. TSS, transcription start site; TES, transcription end site. (D) Western blot of NSD1, GAPDH, and H3K36me2 for WT of NSD1/2-dKO HEK293T cells stably rescued with NSD1 or NSD1^PWWP2–4A^ (E) Neural Progenitor Cell (NPC) differentiation of E14-mESC cells with indicated genotypes. Top, representative images of Embryoid Bodies (EBs) undergoing NPC differentiation after 3 days of Retinoic Acid (RA) treatment. Bottom, quantifications of differentiating and non-differentiating EBs. (F) Western blot of Nestin, GAPDH, and H3K36me2 for E14-mESC cells undergoing EB and NPC differentiation.

### The aromatic pocket of NSD1-PWWP2 is required for NSD1 activation and early stem cell differentiation

Mammalian PWWP domains are comprised of a C-terminal α-helix and a N-terminal β-barrel aromatic pocket that binds to hydrophobic targets^25^. Several PWWP domains were shown to bind methylated lysine of histones, which are considered canonical targets of PWWP^25,26^. In the case of NSD1, the core residues in PWWP2 are RWWP. Since NSD1-PWWP2 does not have any crystal or cryo-EM structures reported in literature, we modeled NSD1-PWWP2 using AlphaFold 2 and compared it with the previously reported NSD3-PWWP2 domain (**Figure S1E**)^27^. The overall architecture of the aromatic pocket of NSD1-PWWP2 is highly conserved with that of NSD3-PWWP2. However, the first core residue R1767 of NSD1-PWWP2 differs in its orientation with its counterpart in NSD3-PWWP2, and we reasoned that this positively charged residue pointing outwards from the aromatic pocket is likely involved in differential target-binding between the two paralogue PWWP domains (**Figure S1E**). W1769 is directly involved in the hydrophobicity of the aromatic pocket and P1770 likely stabilizes the second β-sheet of NSD1-PWWP2 (**Figure S1E**). We reasoned that PWWP2 may be involved in NSD1 allosteric activation by binding external factors through the aromatic pocket. To test this possibility, we generated point mutations in the core residues of NSD1-PWWP2 from RWWP to four alanine (PWWP2-4A) to disrupt the aromatic pocket. Consistent with NSD1^ΔPWWP2^, NSD1^PWWP2–4A^ exhibited little catalytic activity while being expressed in NSD1/2-dKO HEK293T cells (**Figure 1D**), indicating that the aromatic pocket of PWWP2 is required to unleash the catalytic activity of NSD1. Since the NUP98-NSD1 translocation fusion protein has been previously reported as an oncogenic driver and cancer dependency in Acute Myeloid Leukemia (AML), we tested if the aromatic pocket of PWWP2 in NUP98-NSD1 is essential for this oncoprotein^7,22^. Consistent with NSD1^PWWP2–4A^, rescue of NUP98-NSD1^PWWP2–4A^ also exhibited little catalytic activity, underlining PWWP2 as a vulnerability of both WT and NUP98-fused NSD1 (**Figure S1F**). As NSD1 is essential for early ESC differentiation during development^18,21^, we asked if this process also requires the PWWP2-dependent activation of NSD1. We generated a NSD1-KO line in mouse E14Tg2a embryonic stem cells (E14-mESC) by CRISPR/Cas9 (**Figure S1G**). Consistent with the literature, genetic depletion of NSD1 resulted in a global loss of H3K36me2 in ESC (**Figure S1G**). We then rescued NSD1-KO E14-mESC with either WT NSD1 or NSD1^PWWP2–4A^. As expected, re-expression of NSD1 rescued H3K36me2 levels in E14-mESC, but NSD1^PWWP2–4A^ did not (**Figure 1E**). NSD1 mutations in human patients cause Sotos Syndrome, which exhibits prominent neurodevelopmental disorders^16^. Therefore, we adopted a neural progenitor cell (NPC) differentiation assay to interrogate if allosteric regulation of NSD1 by the aromatic pocket of PWWP2 is required for a proper differentiation into neural lineages^28^. We first induced embryoid bodies (EB) formation of E14-mESC and then direct neural lineage differentiation by Retinoic Acid (RA) treatment. EBs are present as spheres in suspension culture and once being induced by RA, they form neuroblasts and migrate outwards from the center of EBs in adherent culture. While both WT and NSD1-KO E14-mESCs can form EBs, NSD1-KO EBs failed to differentiate into the typical neural lineages induced by RA in contrast to WT E14-mESC (**Figure 1F**). Re-expression of WT NSD1 largely rescued this cellular defect in the NSD1-KO background but NSD1^PWWP2–4A^ exhibited little rescue effects (**Figure 1F**). This phenotype is further corroborated by the absence of expression of the early neural differentiation marker, NESTIN, in NSD1-KO and NSD1-KO+NSD1^PWWP2–4A^ cells (**Figure 1E**). These findings demonstrated that NSD1 function in early development is dependent on PWWP2 and strongly implicated that the aromatic pocket of PWWP2 is an allosteric regulation site of NSD1.

### Identification of paraspeckle protein NONO as a non-canonical binding target of NSD1-PWWP2

Having established a critical role of the aromatic pocket of NSD1-PWWP2 in regulating NSD1 activity and ESC differentiation, we next investigated the binding targets of NSD1-PWWP2. Previous work indicated that NSD1-PWWP2 does not bind to NSD1’s own catalytic product, H3K36me2, suggesting that NSD1 does not adopt a “write-and-read” mechanism like PRC2^24^. However, the direct binding targets of NSD1-PWWP2 still remain unclear. We first tested if purified NSD1-PWWP2 and NSD1-PWWP2-4A domains exhibit differential binding to histone PTMs using the MODified^TM^ Histone Peptide Array (Active Motif). However, no significant difference was found (data not shown). We speculated that NSD1-PWWP2 is not a conventional PWWP domain and may bind to non-histone targets. To investigate this possibility, we stably expressed Flag-tagged, full-length WT NSD1 or NSD1^ΔPWWP2^ in both NSD1-KO HEK293T and DIPG13 glioma cells using the PiggyBac system. We then performed Flag-affinity purification and competitive elution using 1X Flag peptide (**Figure 2A**). Eluted NSD1 or NSD1^ΔPWWP2^ from either cell line was subjected to interactomic analyses using liquid chromatography mass spectrometry (LC-MS) (**Figure 2A**). We used a stringent binary cut-off to select candidate PWWP2 interactors that are exclusively present in the interactome of WT NSD1 relative to that of NSD1^PWWP2–4A^ and identified 175 and 42 unique PWWP2-interacting proteins in DIPG13 and HEK293T cells, respectively. Strikingly, Non-POU Domain Octamer-Binding Protein (NONO) appears to be the top candidate in both cell lines with far more peptide-spectrum match (PSM) spectral counts than any other candidates (**Figure 2B**). NONO is a nucleic acid binding protein known to interact with the long non-coding RNA NEAT1 to form nuclear paraspeckles, a membrane-less organelle that retains A-to-I edited mRNAs^29,30^. While NONO has been identified in several IP-MS experiments in literatures as a possible interactor for many nuclear proteins^31–33^, the exact biochemical functions of NONO still remains elusive. The N-terminus of NONO is comprised of two conserved RNA Recognition Motif (RRM) domains, RRM1 and RRM2, which mediate its binding to DNA, RNA, and DNA/RNA hybrids (**Figure 2C**)^34^. The C-terminus of NONO consists of a NONA/paraspeckle (NOPS) domain and a long Coiled Coil domain, which both mediate homo/hetero-dimerization and oligomerization between NONO and its two paralogue proteins, Paraspeckle Component 1 (PSPC1) and Splicing Factor Proline and Glutamine Rich (SFPQ) (**Figure 2C**)^34^. Of note, while SFPQ was not a unique protein identified in our interactomic analysis, it is enriched in the IP of NSD1 relative to that of NSD1^ΔPWWP2^, indicating that a fraction of NONO was bound to SFPQ in the pulldown (**Figure S2A**). In contrast, PSPC1 was not detected in our analyses. To corroborate our IP-MS target discovery results, we first performed a co-IP of full-length NSD1 and NONO and validated that WT NSD1 interacts with NONO while NSD1^ΔPWWP2^ does not (**Figure S2B**). Next, we mapped the interaction between NSD1 and NONO by a GST pulldown assay using GST-NSD1-PWWP2 as the bait. We generated HA-tagged, N-terminus or C-terminus truncated NONO (N-NONO and C-NONO, respectively) and expressed them in HEK293T cells. We found that NSD1-PWWP2 specifically pulls down N-NONO but does not interact with C-NONO, and that such protein-protein interaction is depleted in the pulldown using GST-NSD1-PWWP2-4A (**Figure 2D**). Collectively, we characterized a novel protein-protein interaction between the aromatic pocket of NSD1-PWWP2 and the N-terminus of paraspeckle protein NONO, expanding our current understanding of PWWP domains as merely methylated lysine readers of histones.

**Figure 2.**
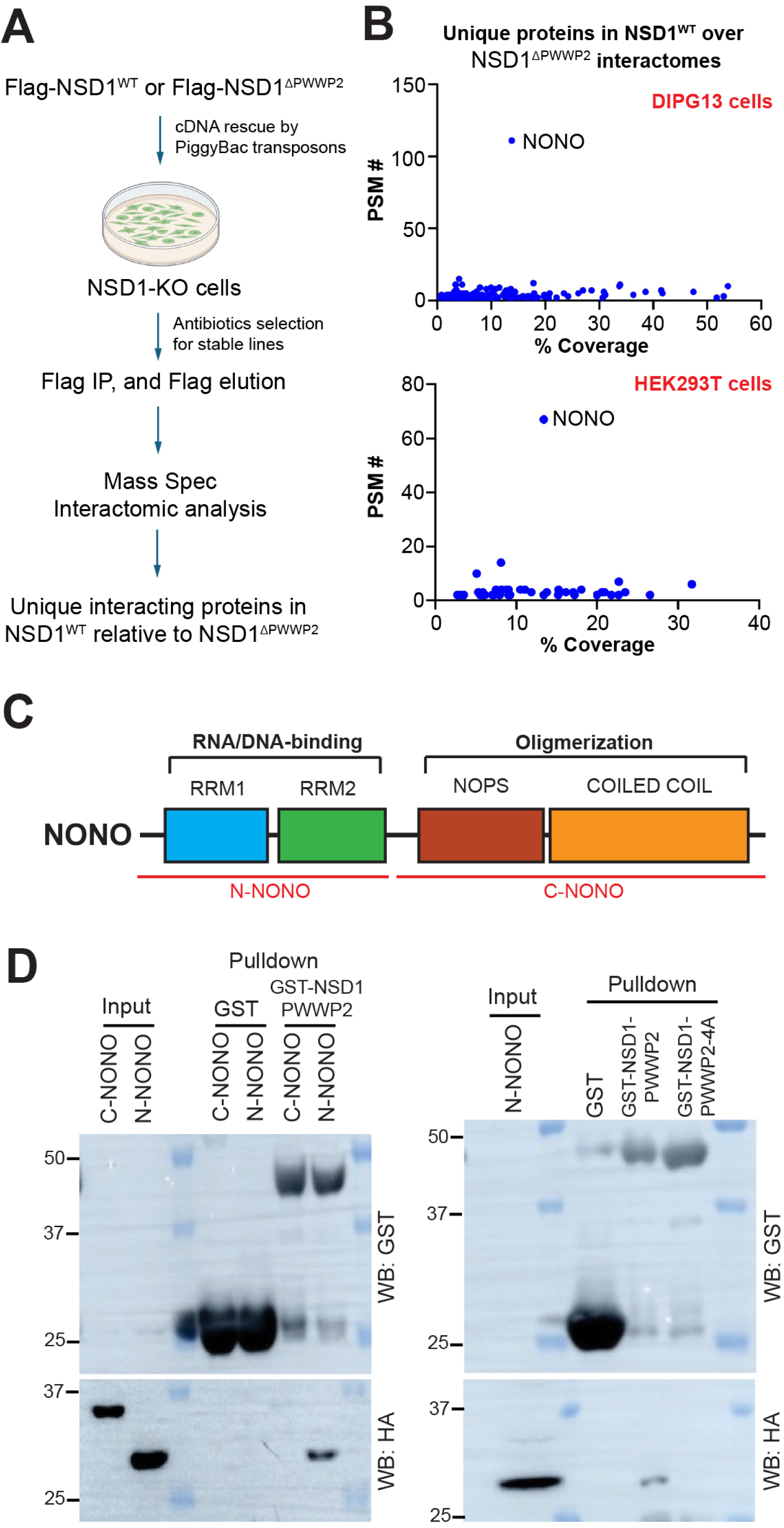
Identification of NONO as a novel NSD1-PWWP2 binding protein through its N-terminal region. (A) Schematic illustration of experimental flows for proteomics analysis of NSD1-PWWP2’s interactomes. (B) Unique proteins detected by LC-MS and plotted by peptide-spectrum match (PSM) scores against percentage of coverage using DIPG13 (top) and HEK293T (bottom) cells. (C) Illustration of annotated functional domains of NONO. (D) GST pulldown assay of HA-tagged NONO using NSD1-PWWP2 as the bait. Left, pulldown of HA-tagged N-NONO or C-NONO using GST alone or GST-NSD1-PWWP2 followed by Western blot of GST and HA. Right, pulldown of HA-tagged N-NONO using GST-NSD1-PWWP2 or GST-NSD1-PWWP2-4A mutant followed by western blot of GST and HA.

### NONO allosterically stimulates NSD1 through the aromatic pocket of PWWP2

We then sought out to characterize the functional role of the NSD1-NONO interaction. As PWWP2 is a critical site for NSD1 activity, we reasoned that NONO may allosterically regulate NSD1 activation. To date, successful biochemical reconstitution of NSD1 for enzymatic assays has been limited to its catalytic module, excluding PWWP2, due to the technical difficulties of reconstituting full-length NSD1 containing substantial unstructured/disordered regions^35^. This hurdle has restricted further investigations into regulatory mechanisms of NSD1. To overcome this limitation, we generated recombinant baculoviruses carrying full-length NSD1 in the Sf9 insect cell expression system and screened for individual monoclonal viruses that stably expressed full-length NSD1 and also produced sufficient protein yield in Sf9 cells (data not shown). With this approach, we obtained clonal viruses that reproducibly express full-length NSD1 and NSD1^PWWP2–4A^ for affinity purification and reconstitution (**Figure 3A**). In addition, we reconstituted N-NONO as well as recombinant di-nucleosomes using the defined 2X Widom 601 sequence as previously described (**Figure 3A**)^36^. We then tested the histone methyltransferase (HMT) activity of full-length NSD1 and NSD1^PWWP2–4A^ on recombinant nucleosomes using a biochemically defined HMT assay. In brief, reconstituted NSD1 and NSD1^PWWP2–4A^ were incubated with recombinant nucleosomes with Tritiated S-Adenosyl methionine ([^3^H]-SAM) as a methyl donor. Covalent incorporation of [^3^H]-methyl groups to H3K36 was detected by autoradiography. With this assay, we demonstrated that an incremental titration of full-length NSD1 exhibited proportionally increased HMT activity in our defined condition (**Figure 3B**). Intriguingly, although full-length NSD1^PWWP2–4A^ had little catalytic activity in rescued HEK293T and E14-mESC cells, it exhibited a similar level of basal HMT activity at the same molarity in comparison with WT NSD1 (**Figure 3B**). We then asked if NSD1 can be allosterically activated by the N-terminus of NONO in the HMT assay. To investigate this, we performed an incremental titration of N-NONO to a fixed molarity of NSD1 and observed that NSD1 catalytic activity is proportionally stimulated by N-NONO (**Figure 3C**). Strikingly, when NSD1^PWWP2–4A^ was subjected to the same experimental condition, it failed to respond to NONO-dependent allosteric stimulation of HMT activity (**Figure 3C**). Together, our results biochemically delineated the basal and allosterically stimulated states of NSD1 and pinpointed the aromatic pocket of PWWP2 as central to NONO-dependent allosteric stimulation of NSD1.

**Figure 3.**
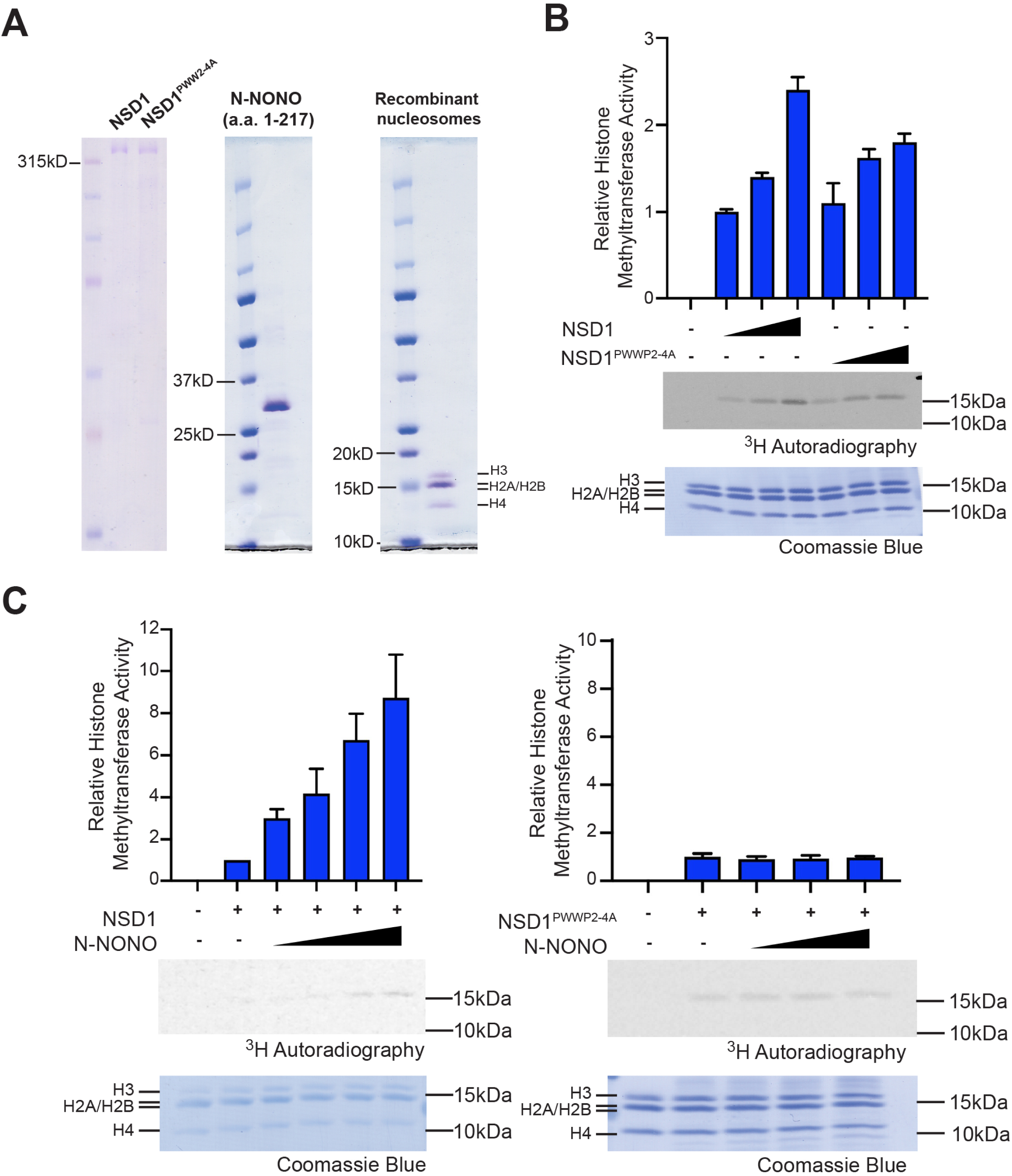
Reconstitution of full-length NSD1 and NSD1^PWWP2–4A^ and identification of NONO as an <u>allosteric activator of NSD1 through the aromatic pocket of PWWP2.</u>. (A) Demonstration of recombinant protein expression and purification including NSD1, NSD1^PWWP2–4A^, N-NONO, and recombinant di-nucleosomes by Coomassie blue staining. (B) HMT assays of full-length NSD1 or NSD1^PWWP2–4A^ mutant in an incremental titration of 62.5, 125, and 250nM. Top, quantifications of autoradiographic signals normalized to NSD1 alone (the second lane). Middle, representative autoradiographic images for stably incorporated [^3^H]. Bottom, Coomassie blue staining of total nucleosomes. (C) HMT assays of 60nM full-length NSD1 with an incremental titration of N-NONO at 0, 530, 880, and 1760nM (left) or 0.25μM NSD1^PWWP2–4A^ mutant with an incremental titration of N-NONO at 0, 880, and 1760nM (right). Top, quantifications of autoradiographic signals normalized to NSD1 alone (the second lane). Middle, representative autoradiographic images for stably incorporated [^3^H]. Bottom, Coomassie blue staining of total nucleosomes.

### NONO/paraspeckle-deficiency dampens steady-state global H3K36me2 deposition

Having uncovered NONO as an allosteric regulator of NSD1, we then asked how this biochemical mechanism impacts NSD1-dependent catalysis of H3K36me2 in cells. To investigate this, we generated NONO-KO E14-mESC cells (**Figure S3A**). We then performed spike-in ChIP-seq experiments for H3K36me2 in steady-state WT and NONO-KO E14-mESC cells. As H3K36me2 spans large active chromatin domains including genic and intergenic regions, we first ranked all regions of the mouse genome from a −10kb of transcription start site (TSS) to +10kb of transcription end site (TES) window based on H3K36me2 enrichment. The meta-analysis profiling and heatmaps of genome-wide H3K36me2 revealed a consistent global reduction of H3K36me2 deposition in both genic and intergenic regions in NONO-KO cells (**Figure 4A and S3B**). Consistently, when we aligned all the broad peaks of H3K36me2 detected in WT and NONO-KO cells, we observed the same global tendency of H3K36me2 reduction on chromatin in NONO-KO cells (**Figure S3B**). As NONO is a core component of nuclear paraspeckles through its binding to lncRNA NEAT1 for oligomerization^29^, we further asked if assemblies of paraspeckles affect NONO-dependent regulation of H3K36me2. We adopted a CRISPR interference (CRISPRi) system by transducing three single-guide RNAs targeting the promoter of NEAT1 along with a dCas9-KRAB construct to knockdown NEAT1 expression. As NEAT1 is not expressed in mESC, we generated NEAT1 CRISPRi HEK293T cells. We validated that CRISPRi resulted in a knockdown of ∼65% of NEAT1 RNA levels by RT-qPCR (**Figure S3C**). We further quantified nuclear paraspeckles by immunofluorescence staining of NONO (**Figure 4B**). We found ∼30% (7/24) of WT cells but only 2.5% (1/40) of NEAT1 CRISPRi cells displayed nuclear paraspeckles, validating our approach to deplete paraspeckle formation (**Figure 4B**). Using H3K36me2 spike-in ChIP-seq and meta-analysis, we found a mild reduction of global H3K36me2 levels in NEAT1 CRISPRi cells compared to WT cells (**Figure 4C and S3D**). However, since paraspeckles are highly dynamic structures observed in only 30% of WT cells, our results demonstrated that paraspeckle assemblies of NONO at least partially affected H3K36me2 deposition. It would be interesting to apply single-cell analysis in the future to compare cells with and without paraspeckles. Overall, these data highlighted a previously unknown role of NONO and paraspeckles in direct regulation of H3K36me2-decorated active chromatin domains.

**Figure 4.**
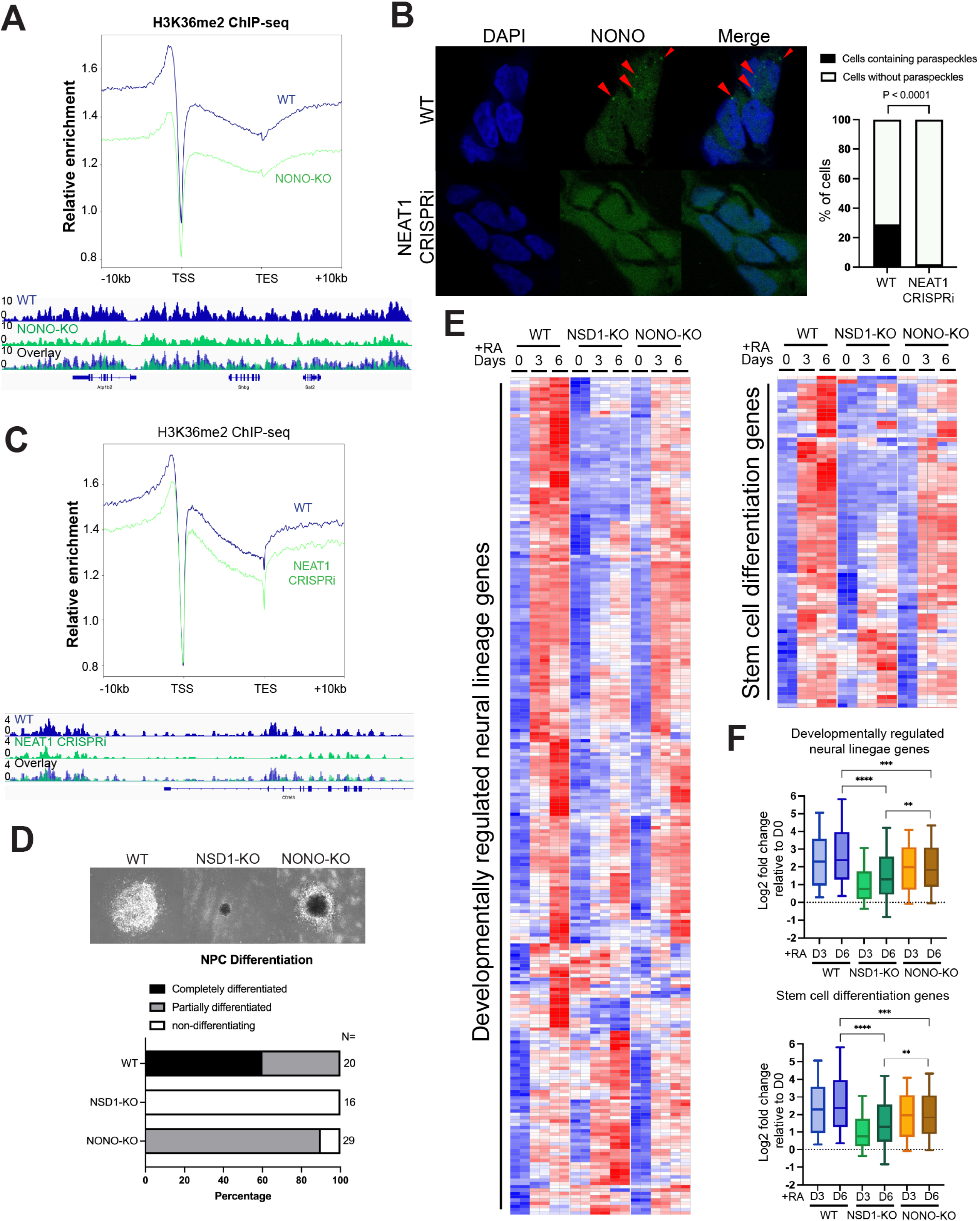
NONO deficiency disrupted global H3K36me2 and partially phenocopied NPC differentiation defects of NSD1-KO. (A) Meta-analysis profiling of H3K36me2 ChIP-seq signals aligned to all genes within a window of −10kb of TSS to +10kb of TES in WT and NONO-KO E14-mESC. Representative track images are shown at the bottom. (B) Left, immunofluorescence staining of NONO in WT and NEAT1 CRISPRi HEK293T cells. Images were captured under a 63X objective and puncta of nuclear paraspeckles were highlighted by red triangles. Right, quantifications of nuclear paraspeckles present in individual WT (n=24) and NEAT1 CRISPRi (n=40) HEK293T cells. The P value is calculated by Chi-squared test. (C) Top, meta-analysis profiling of H3K36me2 ChIP-seq signals in WT and NEAT1 CRISPRi cells aligned to all genes within the same window indicated in (A). Bottom, representative track images of H3K36me2 ChIP-seq in these cells. (D) Neural Progenitor Cell (NPC) differentiation of WT, NSD1-KO, and NONO-KO E14-mESC cells. Top, representative images of Embryoid Bodies (EBs) undergoing NPC differentiation after 3 days of Retinoic Acid (RA) treatment. Bottom, quantifications of fully differentiated, partially differentiated, or non-differentiated EBs. (E) Heatmaps of differential gene expression analysis in WT, NSD1-KO, and NONO-KO cells treated with RA for 0, 3, or 6 days using RNA-seq. 252 genes associated with neural development and 102 genes associated with stem cell differentiation were presented. (F) Box plots of log2 fold changes in gene expression using the experimental conditions shown in (D). The box and whisker represent 95%, third quartile, median, first quartile, and 5% distribution of genes. P values were calculated by Wilcoxon test **p<0.01; ***p<0.001; ****p<0.0001.

### NONO deficiency partially phenocopies the neural differentiation defect of NSD1 knockout

NONO deficiency has been previously reported to cause Syndromic X-linked Intellectual Disability 34 (MRXS34), also called Macrocephaly-intellectual disability-left ventricular non-compaction syndrome, in male patients^37^. Intriguingly, both Sotos Syndrome (NSD1 mutant) and MRXS34 Syndrome patients exhibited macrocephaly, a rare neurodevelopmental phenotype with little mechanistic understanding^37^. In parallel to our findings on the biochemical regulation of NSD1 by NONO, this genetic correlation between NSD1 and NONO in human brain disorders urged us to further investigate the cellular phenotypes of NONO deficiency in early neural differentiation. To investigate this, we first tested the differentiation potential of NONO-KO E14-mESC in comparison with WT and NSD1-KO cells using the NPC differentiation assay. We classified EBs into three categories, fully differentiated, partially differentiated, and non-differentiating based on their morphology since differentiated NPCs manifest a flattened, adherent cellular morphology in 2D culture in contrast to EBs that manifest sphere structures. As 100% (20/20) of WT and 0% (0/16) of NSD1-KO EBs are fully or partially differentiated by the induction with RA, NONO-KO EBs manifested an intermediate phenotype, with 0% (0/29) undergoing full differentiation, 90% (26/29) partial differentiation and 10% (3/29) no differentiation (**Figure 4D**). Next, we characterized the differential gene expression underlying this phenotype of the impaired neural differentiation of NONO-KO E14-mESC in comparison with WT and NSD1-KO cells by RNA-seq. We adopted Gene Ontology for Biological Process gene signatures and examined 252 genes associated with neural differentiation, central nervous system development, and neurogenesis, as well as 102 genes associated with stem cell differentiation (**Supplemental Table 1 and 2**)^38^. We generated heatmaps for all developmentally regulated neural lineage genes and stem cell differentiation genes as well as clustering of gene signatures to compare the changes of WT, NSD1-KO, and NONO-KO cells in response to RA-induced NPC differentiation (**Figure 4E, S3E, and Supplemental Table 1 and 2**). While NSD1-KO cells showed a drastic downregulation of these genes and signatures compared to WT cells, NONO-KO cells displayed an intermediate decrease, consistent with the cellular phenotype in the NPC differentiation assay (**Figure 4E and S3E**). To quantitatively analyze differential gene expression associated with the NPC differentiation phenotype, we plotted the log2-scale fold change in gene expression for Day 3 and Day 6 of RA-induction normalized to the Day 0 baseline expression levels for all three genotypes (**Figure 4F**). We observed significant differential gene expression for each comparison between WT, NSD1-KO, and NONO-KO cells undergoing NPC differentiation and corroborated that NONO-KO manifested a partially impaired differentiation phenotype relative to NSD1-KO (**Figure 4F**). These data characterized the cellular and transcriptional defects in NPC differentiation of NSD1-KO and NONO-KO cells, demonstrating how altered H3K36me2 levels determined phenotypic severity and providing a molecular basis of understanding chromatin regulation defects underlying NONO-deficiency.

## DISCUSSION

Since H3K27 and H3K9 methyltransferases possess an intrinsic “write-and-read” mechanism, the prevailing model of how a balance between active and repressive chromatin is established argues that repression is the default of chromatin state and that active chromatin histone PTMs are “passively” deposited following active transcription^39,40^. However, this model does not fully explain how H3K36me2 domains are placed spanning mega-base long intergenic regions devoid of active transcription. Since the initial discovery of NSD1 methyltransferase activity, the biochemical and structural characterizations have been largely restricted to the catalytic module. In this study, we overcome the technical hurdles to establish the first biochemically defined HMT assay for full-length NSD1, allowing us to interrogate how NONO allosterically regulates its activity. Our findings on the NONO-NSD1 regulation shed new lights into the molecular underpinnings of how H3K36me2 domains are established and underlined that an allosteric mechanism of NSD1 independent of transcription should be considered to revise the current model of chromatin balance between active and repressive domains. As NONO was reported to be physically associated with active chromatin including intergenic regions^31^, it would be an interesting future direction to explore how NONO is temporo-spatially placed on chromatin to regulate NSD1 activity to ensure faithful inheritance of H3K36me2 domains after DNA replication. Although NONO has been shown to interact with several protein machineries and serves as a key structural component of nuclear paraspeckles^31–33^, the definitive, biochemical role of NONO and paraspeckles remained largely elusive. Our study uncovered a direct mechanistic link between NONO/paraspeckles and NSD1 and elucidated a novel function of NONO/paraspeckles despite its many proposed roles.

Another surprising finding here is the target-binding specificity of NSD1-PWWP2. To our knowledge, mammalian PWWP domains were only known to bind methylated lysine of histones for chromatin signaling until a recent study demonstrating that a C-terminal fragment of NSD1 binds to H3K18ac and suggested that the PHD1 to PHD4 domains and PWWP2 are simultaneously involved^21,25,26^. However, our data showed that PWWP2 alone is sufficient to bind to NONO through its aromatic pocket, characterizing the first non-histone binding target of mammalian PWWP domains. Our finding on the NSD1-PWWP2-NONO interaction along with the recent report demonstrated the possible target-binding diversity of mammalian PWWP domains that may bridge unexpected crosstalk between distinct chromatin-regulating machineries. In addition, our results also highlighted the aromatic pocket of PWWP2 as a vulnerability of NSD1 for cancer therapy. Currently, there are no potent, clinical-grade inhibitors targeting the SET domain of NSD1 for clinical trials against NSD1-dependent cancer, such as DMG or NUP98-NSD1 fusion AML. Our study suggests that PWWP2 may serve as a pharmacological target for the development of allosteric inhibitors against NSD1 to circumvent current hurdles to develop orthosteric inhibitors.

While NSD1^PWWP-4A^ lacking allosteric activation failed to rescue the NPC differentiation defect in NSD1-KO E14-mESC, NONO-KO only exhibited a partial defect. We reason that the loss of NONO does not fully phenocopy the loss of NSD1 allosteric activation because there likely are additional allosteric activators in parallel to NONO. This observation is reminiscent of JARID2, a co-factor of PRC2 that also allosterically stimulates PRC2 in parallel to H3K27me3^41–43^. While PRC2 null ESC failed to differentiate in any of the three germ layers, JARID2-KO ESC selectively manifested a developmental defect of mesodermal genes^44^. Our successful reconstitution of full-length, catalytically active NSD1 provided a new platform for future characterization of additional allosteric regulators of NSD1.

While NSD1-KO is embryonic lethal, heterozygous NSD1 mutations cause human Sotos Syndrome that manifests overgrowth in multiple types of tissues including macrocephaly of the brain^16^. Intriguingly, MRXS34 Syndrome caused by hemizygous X-linked NONO mutations in male only exhibits macrocephaly but not overgrowth in other organs^37^. Based on our data, a partially impaired neural differentiation may result in aberrant accumulation of stem cell pools and expansion in brain overgrowth. We speculate that NONO might be the major allosteric activator of NSD1 in the ectodermal and neural lineages. Future investigations are certainly warranted to identify lineage-specific allosteric regulators of NSD1 and H3K36me2-decorated active chromatin.

Overall, we provided critical advances to the biochemical basis of active chromatin formation and its relationship with nuclear paraspeckles. Our work further implicated a new pharmacological target site in NSD1 for therapeutic interventions and a molecular insight towards understanding two poorly studied human neurodevelopmental disorders, Sotos and MRXS34 Syndromes.

## MATERIAL AND METHODS

### cDNAs and Expression vector

For recombinant protein expression in Sf9 cells, human NSD1 cDNA (GeneScript, catalog no. OHu18754) was cloned to pFASTBac1 with 1x Flag tag at the N-terminus end. The ΔRBD1, ΔRBD2/3, ΔPWWP1, and ΔPWWP2 mutants were then generated by site-directed mutagenesis (Agilent catalog no. 200523). The cDNAs of NSD1-PWWP2 (a. a. 1695-1873) fragments were amplified from NSD1-WT or NSD1-PWWP2-4A by PCR and cloned to pGEX4T2 for expressing in bacteria. cDNAs of full-length, a. a. 1-217, and a.a. 218-466, of human NONO were amplified from NONO cDNA (SinoBiological, catalog no. HG14826-CF) and cloned to pFASTBac1 with 1x Flag tag at the N-terminus end. CL20_mEGFP_NUP98_NSD1 was purchased from Addgene (# 205873) and the PWWP2-4A mutant was generated by site-directed mutagenesis. The cDNAs described above were sub-cloned to pPB-CAG-3xFLAG-empty-pgk-hph (Addgene # 48754) or pcDNA3.1(+) (Invitrogen) according to experimental needs.

### Antibodies

The following antibodies were used for western blot and ChIP-seq: NSD1 (University of California, Davis/National Institutes of Health NeuroMab Facility) mouse monoclonal, N312/10; NSD1 (Abbexa) rabbit polyclonal, catalog no. abx135901; GAPDH (ProteinTech) mouse monoclonal 1E6D9, catalog no. 60004-1-Ig; Nestin (Cell Signlaing Technology) mouse monoclonal 10C2, catalog no. 33475; Anti-HA-tag (Cell Signaling Technology) rabbit monoclonal C29F4, catalog no. 3724; H3K36me2 (Cell Signaling Technology) rabbit monoclonal C75H12, catalog no. 2901; H3K36me2 (Cell Signaling Technology) rabbit monoclonal C75H12 (ChIP Formulated), catalog no. 33587; NSD2 (Millipore) mouse monoclonal 29D1, catalog no. MABE191; NONO (ProteinTech) rabbit polyclonal, catalog no. 11058-1-AP; NONO (Santa Cruz Biotechnology) mouse monoclonal A-11, catalog no. sc-166702; histone H3 (Abcam) rabbit monoclonal EPR16987, catalog no. ab176842; anti-Flag (Sigma-Aldrich) mouse monoclonal M2, catalog no. F1804; HRP-conjugated GST Tag (ProteinTech) mouse monoclonal antibody, catalog no. HRP-66001 and H2Av (Active Motif) rabbit polyclonal, catalog no. 39715.

### ChIP-seq and RNA-seq

ChIP-seq experiments were performed as previously described^36^. Briefly, cells were cross-linked with 1% formaldehyde for 10 min. Following nuclei isolation, the chromatin was extracted and fragmented to ∼250 base pairs (bp) using a Diagenode Bioruptor. ChIP was performed with the specific antibodies listed above. For quantification, (spike-in) chromatin from Drosophila (1:100 ratio to the experimental chromatin) with Drosophila-specific H2Av antibody was added to each sample as a spike-in control, allowing ChIPs to be compared to one another. Libraries were prepared using 1 to 30 ng of immunoprecipitated DNA. For RNA-seq experiments, RNA was isolated using GeneJet RNA purification kit (Thermo Scientific) according to the manufacturer’s instruction. Libraries were constructed using NEB Next Ultra II library preparation kit. Libraries were sequenced at 40M paired-end reads for each sample (AmpSeq).

### ChIP-seq data analysis

150bp paired-end reads were generated by AVITI sequencing (Element Biosciences). ChIP-seq analysis was performed with the nf-core/chip-seq v2.1.0 pipeline using the Nextflow-based nf-core framework in a singularity container^45–48^. Briefly, reads were pre-processed with cutadapt5 and FastQC [https://www.bioinformatics.babraham.ac.uk/projects/fastqc/] aligned to either hg38 or dm6 genomes with bowtie2, and reads were further cleaned and sorted with Picard [http://broadinstitute.github.io/picard/] and SAMtools^49,50^. Using the number of uniquely mapped paired reads to either the hg38 or dm6 genomes, the ratio of hg38 to dm6 reads were calculated by dividing hg38 reads by dm6 reads. The wild type sample’s ratio was set to 1, and mutant samples were calculated relative to the wild type. Coverage tracks were then generated using the bam_coverage function of deepTools, with the scaling factor manually set as calculated for each sample^51^. Tracks were then visualized in the IGV browser^52^. Heatmaps and metagene plots were created with the deepTools suite. Pipeline outputs and parameters can be found at (data upload location). Raw and processed data are available at (GEO Reference).

### RNAseq data processing and GSEA analysis

Sequencing reads were aligned to mm10 reference genome using STAR v2.7.6a [https://pmc.ncbi.nlm.nih.gov/articles/PMC3530905/]. Gene-level quantification was carried out using featureCounts v2.0.0 to generate a raw counts matrix^53^. To calculate normalized expression levels, FPKM (Fragments Per Kilobase of transcript per Million mapped reads) values were estimated using Cufflinks v1.46^54^. Differential gene expression analysis was conducted using DESeq2 v3.20 with default parameters^55^. Gene Set Enrichment Analysis (GSEA) was performed using the clusterProfiler v2.11 R package^56^. A ranked gene list was generated based on z-scores, and the analysis was performed on C5:GO:BP biological pathway categories. Statistically significant genes were determined by p-adjust value 0.05.

### Immunofluorescence staining

HEK293T cells were grown on poly-D-lysine coated cover slides and fixed with 4% paraformaldehyde (Sigma) for 10 min at room temperature. Cells were penetrated and blocked with blocking buffer (1x PBS, 1% BSA, 0.3% Triton-x 100, 0.02% NaN3) at room temperature for 1 hour. 5ng/µL of primary antibody was diluted in antibody dilution buffer (1x PBS, 1% BSA, 0.02% NaN3) and incubate with cells at 4°C overnight. Fluorochrome-conjugated secondary antibody (Alexa Fluor 488) was diluted to 10pg/µL with antibody dilution buffer and incubate with cells at room temperature for 1 hour. The coverslip and the stained cells were mount with mounting medium (Invitrogen). The cover slide and cells were washed three times with PBS between each of the above steps.

### Production of polyclonal baculovirus

Flag-NSD1-WT, Flag-NSD1-PWWP2-4A, Flag-NONO, Flag-N-NONO (aa1-217) were cloned into pFASTBac1 (Invitrogen) and used to produce P0 polyclonal baculovirus according to Invitrogen guideline for the Bac-to-Bac system. Sf9 cells was infected with P0 polyclonal baculovirus to produce P1 polyclonal baculovirus, and so on. P2 and P3 polyclonal baculovirus were used for expression of Flag-NONO and Flag-N-NONO for protein purification.

### Isolation of monoclonal baculovirus

A large number of non-recombinant viruses were selected against large protein expression, such as full-length NSD1, using the Bac-to-Bac system. To overcome this issue, we performed a plaque assay to isolate monoclonal baculovirus. In brief, P0 polyclonal baculovirus with Flag-NSD1-WT or Flag-NSD1-PWW2-4A inserts were prepared according to Invitrogen guidelines. One million per well of Sf9 cells were placed in a 6-well plate, serially diluted P0 polyclonal virus was added to each well, and incubated at 27.5°C for 3 h. All the culture media in the wells was removed, and 3mL of top agarose (ESF-921 at 42°C with microwave-boiled 3% (w/v) agarose in a 9:1 ratio; mix and use immediately) was rapidly added to each well. The 6-well plate was incubated at room temperature until top agarose solidified, then incubated at 27.5°C for 2 weeks. After the plaques were enlarged to a visible size, the top agarose containing the virus was picked up from the single plaque with a P2 tip and transferred into a 12-well plate pre-seeded with 0.5 million per well of Sf9 cells. Incubate the infected Sf9 cells at 27.5°C until >50% of cells burst, then harvest P0 monoclonal baculovirus. Infect Sf9 cells with a portion of the P0 baculovirus and perform Western blotting to compare the protein expression level of each clone. The high-performed monoclonal baculovirus were amplified according to Invitrogen’s guidelines.

### Protein purification

NSD1 and N-NONO were expressed separately in Sf9 by baculovirus infection. After 72 hours of infection, Sf9 cells were suspended lysis buffer (20mM HEPES-NaCl pH7.8, 1mM EDTA, 350mM NaCl, 10% (v/v) glycerol, 0.1% (v/v) NP-40 alternative) with protease inhibitors (1 mM phenylmethlysulfonyl fluoride (PMSF), 0.1 mM benzamidine, 1.25 mg/mL leupeptin 1mg/mL aprotonin and 0.625 mg/mL pepstatin A). Cells were lysed by sonication (Fisher Sonic Dismembrator model 120), and NSD1 or N-NONO were affinity purified through FLAG-M2 agarose beads (Sigma) and eluted with 100µg/mL of 1x Flag-peptide (MedChemExpress). Eluted protein was dialyzed in storage buffer (20mM HEPES-NaCl pH7.8, 1mM EDTA, 350mM NaCl, 5% (v/v) glycerol, 0.02% (v/v) NP-40 alternative). Proteins were concentrated with Amicon® Ultra Centrifugal Filter (Sigma) after dialysis and then snap-frozen in liquid nitrogen and stored at −80°C. A small amount of the sample is retained at each step for analysis of yield, purity and concentration.

### HMT assay

Standard HMT assays were performed in a total volume of 25 μL containing HMT buffer (50 mM Tris-HCl, pH 8.5, 5 mM MgCl2, and 4 mM DTT) with 0.5 µM of [^3^H]-labeled S-Adenosylmethionine (SAM, Perkin Elmer), 60 nM (500 ng) of recombinant dinucleosomes consisting of 2x Widom 601 sequence (120 nM of nucleosome), recombinant human NSD1s, and N-NONO. The reaction mixture was incubated for 90 min at 30°C and stopped by the addition of 6.5 μL SDS buffer (0.2 M Tris-HCl, pH 6.8, 20% glycerol, 10% SDS, 10 mM β-mercaptoethanol, and 0.05% Bromophenol blue). A titration of NSD1 was performed under these conditions to optimize the HMT reaction within a linear range, and the yield of each HMT reaction was measured using the following procedures. After HMT reactions, samples were incubated for 5 min at 95°C with gel loading buffer and separated on SDS-PAGE gels. The gels were then subjected to wet transfer of proteins to 0.45 μm PVDF membranes (Millipore). The radioactive signals were detected by exposure on autoradiography films (Denville Scientific). All HMT assays were performed with the above indicated concentrations of NSD1 and N-NONO unless otherwise noted in the figure legends.

## CONTRIBUTIONS

J-R.Y. conceived and supervised the study and wrote the manuscript. C-I.H and J-R.Y. conducted most of the experiments. S.M. and J.D. analyzed deep-sequencing data. S.R. assisted with stem cell differentiation experiments. M.S. assisted with clonal selection of cell lines.

## ACKNOWLEDGEMENTS

We thank past and current Yu Lab members for technical supports. We also thank D. Reinberg (U Miami/HHMI) for generously providing critical reagents, C-H. Lee (Seoul National) and C. Chen (UBC) for constructive feedback, and B. Ueberheide and the NYU proteomics core for LC-MS analysis. J-R.Y. was supported by Virginia Tech start up fund, Virginia Tech Seale Innovation Fund, CURE Childhood Cancer Fund, Children’s Cancer Research Fund, and The Children’s Cancer Foundation. S.M. was supported by Prostate Cancer Foundation Young Investigator Award.

## DECLARE OF INTEREST

The authors do not have any conflict of interest to declare.

## Notes

### Competing Interest Statement

The authors have declared no competing interest.

