## Supplemental Figures for "Paraspeckle Protein NONO Regulates Active Chromatin by Allosterically Stimulating NSD1"

**Supplemental Information**

**Figure S1 (related to Figure 1)**

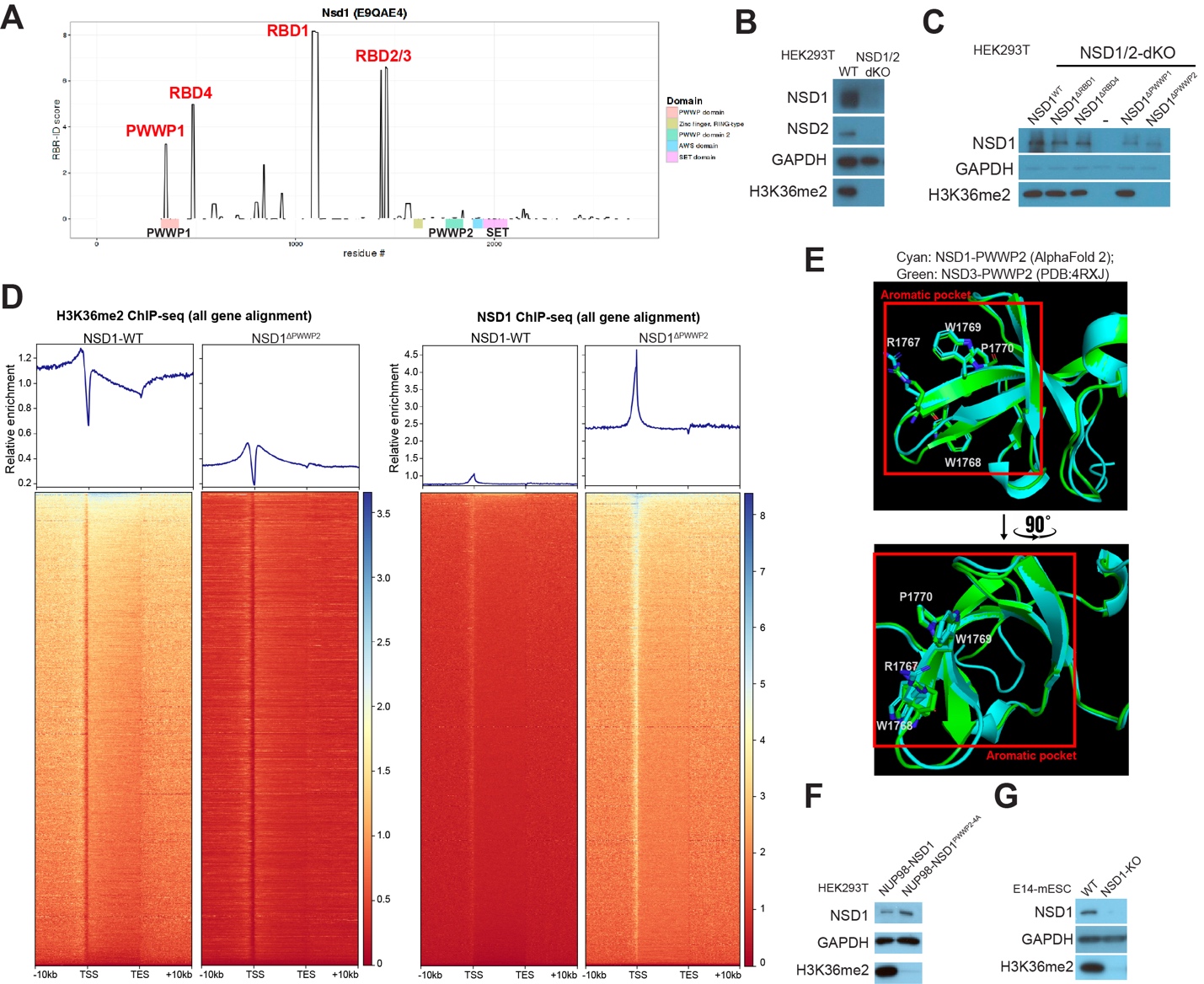

(A) Schematic illustration of RNA-binding domains (RBD) detected by RBR-ID
(B) Western blot of NSD1, NSD2, GAPDH, and H3K36me2 in WT and NSD1/2-dKO HEK293T cells.
(C) Western blot of NSD1, GAPDH, and H3K36me2 in NSD1/2-dKO HEK293T cells stably rescued with WT or mutant NSD1.
(D) Meta-analysis profiling and heatmaps of H3K36me2 ChIP-seq (left) and NSD1 ChIP-seq (right) aligned to all genes within -10kb of TSS to +10kb of TES.
(E) AlphaFold2-modeled aromatic pocket of NSD1-PWWP2 (cyan) overlayed with that of NSD3-PWWP2 (green, PDB: 4RXJ). The core residues of NSD1-PWWP2 were displayed with side chains (R1767, W1767, W169, and P1770).
(F) Western blot of NSD1, GAPDH, and H3K36me2 of NSD1/2-dKO rescued with NUP98-NSD1 or NUP98-NSD1^PWWP2-4A^ in HEK293T cells.
(G) Western blot of NSD1, GAPDH, and H3K36me2 of WT and NSD1-KO E14-mESC.

**Figure S2 (related to Figure 2)**

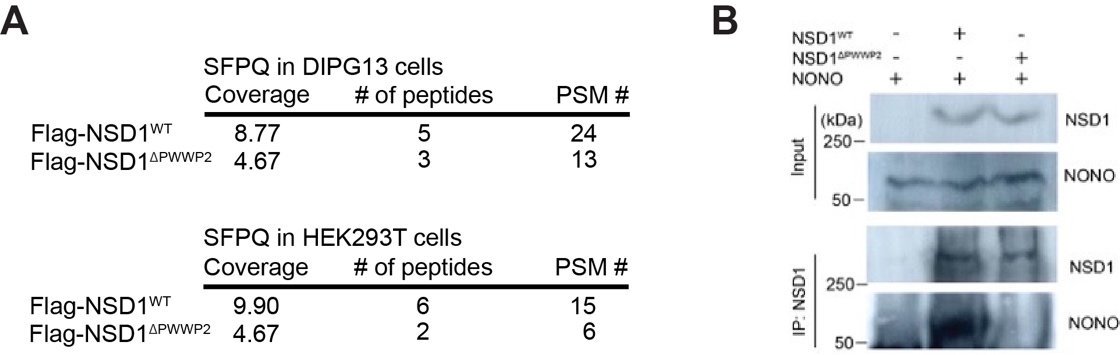

(A) Detection of SFPQ in the interactomic analysis by LC-MS in DIPG13 and HEK293T cells.
(B) Co-Immunoprecipitation (Co-IP) using Flag-tagged NSD1 or NSD1ΔPWWP2 as the bait for the pulldown of NONO, followed by western blot using NSD1 and NONO antibodies.

**Figure S3 (related to Figure 4)**

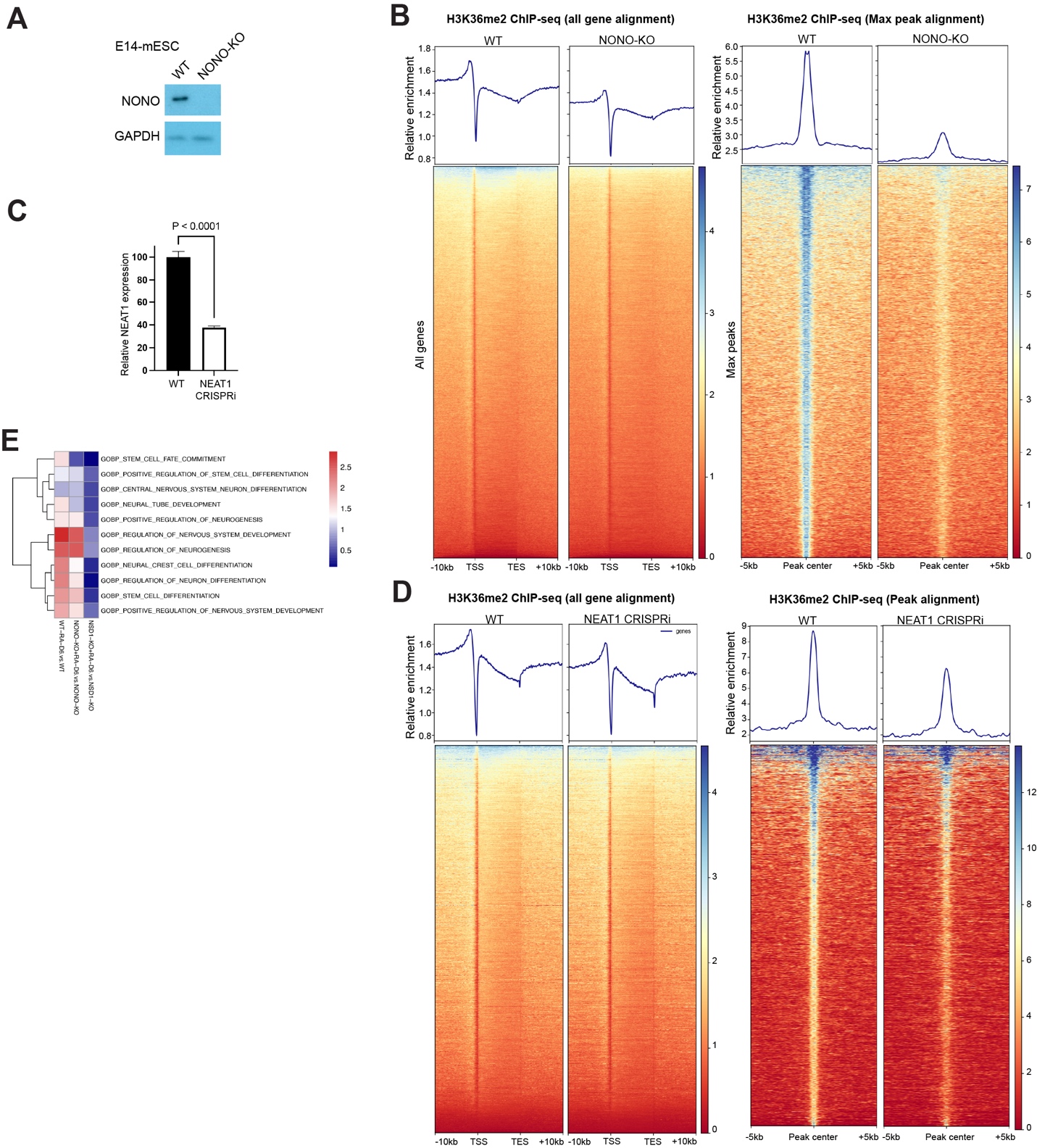

(A) Western blot of NONO and GAPDH in WT and NONO-KO E14-mESC
(B) Meta-analysis profiling and heat maps of H3K36me2 ChIP-seq in WT and NONO-KO E14-mESC cells. ChIP-seq signals were aligned to all genes within a -10kb of TSS to +10kb of TES window or to peak center.
(C) qPCR quantification of NEAT1 RNA expression levels in WT and NEAT1 CRISPRi cells. Signals were normalized by GAPDH. N=5 for each condition. P value was calculated by student’s t-test.
(D) Meta-analysis profiling and heat maps of H3K36me2 ChIP-seq in WT and NEAT1 CRISPRi HEK293T cells. ChIP-seq signals were alighed to all genes within a -10kb of TSS to +10kb of TES window.
(E) Heatmap of significantly changes of Gene Set Enrichment Analysis (GSEA) signatures including stem cell differentiation and neural lineage gene sets in WT compared to NSD1-KO, and NONO-KO E14-mESC cells undergoing RA induced NPC differentiation.
